# Evidence for reduced long-term potentiation-like visual cortical plasticity in schizophrenia and bipolar disorder

**DOI:** 10.1101/2020.06.06.128926

**Authors:** Mathias Valstad, Daniël Roelfs, Nora B. Slapø, Clara M.F. Timpe, Ahsan Rai, Anna Maria Matziorinis, Dani Beck, Geneviève Richard, Linn Sofie Sæther, Beathe Haatveit, Jan Egil Nordvik, Christoffer Hatlestad-Hall, Gaute T. Einevoll, Tuomo Mäki-Marttunen, Marit Haram, Torill Ueland, Trine V. Lagerberg, Nils Eiel Steen, Ingrid Melle, Lars T. Westlye, Erik G. Jönsson, Ole A. Andreassen, Torgeir Moberget, Torbjørn Elvsåshagen

## Abstract

**Background:** Several lines of research suggest that impairments in long-term potentiation (LTP)-like synaptic plasticity might be a key pathophysiological mechanism in schizophrenia (SZ) and bipolar disorder type I (BDI) and II (BDII). Using modulations of visually evoked potentials (VEP) of the electroencephalogram, impaired LTP-like visual cortical plasticity has been implicated in patients with BDII, while there has been conflicting evidence in SZ, a lack of research in BDI, and mixed results regarding associations with symptom severity, mood states, and medication.

**Methods:** We measured the VEP of patients with SZ spectrum disorders (n=31), BDI (n=34), BDII (n=33), and other BD spectrum disorders (n=2), and age-matched healthy control participants (n=200) before and after prolonged visual stimulation.

**Results:** Compared to healthy controls, modulation of VEP component N1b, but not C1 or P1, was impaired both in patients within the SZ spectrum (χ^2^=35.1, p=3.1×10^−9^) and BD spectrum (χ^2^=7.0, p=8.2×10^−3^), including BDI (χ^2^=6.4, p=0.012), but not BDII (χ^2^=2.2, p=0.14). N1b modulation was also more severely impaired in SZ spectrum than BD spectrum patients (χ^2^=14.2, p=1.7×10^−4^). The reduction in N1b modulation was related to PANSS total scores (χ^2^=10.8, p=1.0×10^−3^), and nominally to number of psychotic episodes (χ^2^=4.9, p=0.027). *Conclusions.* These results suggest that LTP-like plasticity is impaired in SZ and BDI, but not BDII, and related to psychotic symptom severity. Adding to previous genetic, pharmacological, and anatomical evidence, these results implicate aberrant synaptic plasticity as a mechanism underlying SZ and BD.

## Introduction

Schizophrenia (SZ) and bipolar disorders (BD) are severe psychiatric disorders, with a lifetime prevalence of ∼0.7% (1) and ∼2% (2,3), respectively. While their precise neural substrates remain unknown despite decades of research, recent genetic, pharmacological, and imaging evidence has implicated aberrant synaptic plasticity as a leading candidate mechanism in SZ and BD (4-6).

Through genome-wide association studies, increased risk of SZ and BD have been associated with single nucleotide polymorphisms (SNPs) at genes that are directly or indirectly involved in glutamatergic synaptic plasticity, such as *GRIN2A* and *CACNA1C* (5,7-9). Moreover, medical conditions affecting N-methyl-D-aspartate (NMDAR) function, including systemic lupus erythematosus (4) and anti-NMDAR encephalitis (5), are associated with distinct psychotic symptoms. Negative symptoms as well as hallucinations, both characteristic of SZ, are reliably produced by NMDAR antagonists such as phencyclidine (6) and ketamine (7), further suggesting that aberrations in NMDAR-dependent synaptic function, and likely in synaptic plasticity in particular (8), constitute a key pathophysiological mechanism in psychotic disorders.

A well-characterized non-invasive marker for NMDAR-dependent LTP-like visual cortical plasticity can be obtained in humans and other species by using EEG to measure modulations of VEP after high-frequency or prolonged visual stimulation (9-11). In rodents, NMDAR antagonists as well as α-amino-3-hydroxy-5-methyl-4-isoxazolepropionic acid receptor (AMPAR) insertion-inhibitor GluR1-CT prevent VEP modulation (12). Further, ζ inhibitory peptide, an inhibitor of Protein Kinase M ζ, which is crucial for maintenance of LTP, disrupts the retention of VEP modulation (13). Further, electric tetanus-induced LTP in the primary visual cortex modulates VEP, and inhibits further visual stimulation-induced VEP modulation (14). Such results strongly suggest that visual stimulation-induced VEP modulation and LTP share common underlying mechanisms.

With the VEP modulation paradigm, aberrant LTP-like plasticity has been implicated in BDII (15,16), and in major depression (MDD) (9). However, there is no previous study of VEP modulation in BDI, and results in SZ have been inconsistent (17,18), possibly due to differences in the visual stimulation applied (18). Efforts have been made to associate VEP modulation with symptom severity, mood states, and medication in order to assess the trait stability of VEP modulation impairments, with mixed results (9,15,16). Thus, further evidence is required to establish impaired LTP-like synaptic plasticity as a disease characteristic in psychotic disorders.

Here, we compared patients with SZ, BDI, or BDII, and healthy controls with respect to VEP modulation after prolonged visual stimulation, with the primary aim to examine whether LTP-like visual cortical plasticity is affected in these disorders. Further, our secondary aims were to i) examine the pairwise differences in VEP modulation between diagnoses, ii) examine the association between VEP modulation and illness severity in patients, and iii) examine the association between VEP modulation and current use of psychotropic medications.

## Methods and materials

### Participants

One hundred patients with BD type I (n=34), BD type II (n=33), SZ (n=25), schizophreniform disorder (n=3), schizoaffective disorder (n=3), BD not otherwise specified (NOS) (n=1), and cyclothymia (n=1), and 411 healthy volunteers were included in this study. Since patients’ ages ranged from 18 to 69 years, while ages of healthy controls ranged from 20 to 90 years, with means of 36.1 and 49.0 years, respectively, all analyses were performed using an age-matched healthy control sample (n = 200, age range: 20 to 70, mean age: 38.2), drawn using the nearest neighbor method in the `MatchIt` package in R (19). Patients were recruited through psychiatric in- and outpatient treatment units in the Oslo area, while healthy controls were recruited through national records and advertisements in a regional newspaper (20). Participants with known neurological disorders, in addition to those who had been subjected to moderate to severe head injury at any time in their life, were excluded from the study. All participants had normal or corrected-to-normal vision. The study was approved by the Regional Ethical Committee of South-Eastern Norway, and all participants provided written informed consent before the experiment started. A detailed characterization of the sample is provided in Table 1.

**Table 1.** Values represent either number of participants, or mean and standard deviation. **HC**: Healthy controls. **SZ**: Schizophrenia spectrum disorders. **BD**: Bipolar spectum disorders. **p**: Probability of no difference between the three groups. **MADRS**: Montgomery and Asberg Depression Rating Scale. **YMRS**: Young Mania Rating Scale. **PANSS**: Positive and Negative Symptoms Scale. **IDS**: Inventory of Depressive Symptoms. **T/L/V/Pr/U**: Total antiepileptics / Lamotrigine / Valproate / Pregabaline / Unspecified.

|  | HC (n = 190) | SZ (n = 29) | BD (n = 68) | p |
| --- | --- | --- | --- | --- |
| Age | 37.3 (11.1) | 36.7 (11.2) | 36.0 (12.6) | 0.24 |
| Sex (f/m) | 107/83 | 13/16 | 46/22 | 0.09 |
| IQ | 114.8 (10.8) | 104.7 (13.9) | 114.6 (11.9) | $8.3 \times 10^{-5}$ |
| Illness duration (years) | – | 11.0 (9.5) | 10.4 (10.2) | 0.81 |
| MADRS | 1.7 (2.6) | 12.3 (6.4) | 14.6 (9.6) | $6.5 \times 10^{-67}$ |
| YMRS | 0.8 (1.5) | 3.8 (4.8) | 4.1 (5.0) | $8.2 \times 10^{-15}$ |
| PANSS total | – | 61.3 (13.7) | 43.7 (8.4) | $4.8 \times 10^{-14}$ |
| IDS sleep items | 1.6 (1.8) | 3.9 (2.2) | 3.8 (2.5) | $6.3 \times 10^{-17}$ |
| No psychotropics | 190 | 5 | 15 | $5.6 \times 10^{-45}$ |
| Antipsychotics | 0 | 20 | 24 | $1.1 \times 10^{-26}$ |
| Antiepileptics (T/L/V/Pr/U) | 0 | 1/0/1/0/0 | 24/18/3/1/2 | $5.3 \times 10^{-18}$ |
| Antidepressants | 0 | 4 | 28 | $2.2 \times 10^{-19}$ |
| Anxiolytics/hypnotics | 0 | 2 | 6 | $2.7 \times 10^{-4}$ |
| Lithium | 0 | 1 | 9 | $2.2 \times 10^{-6}$ |
| Tobacco daily (y/n) | 40/141 | 12/16 | 27/39 | 0.003 |
| Cannabis last month (y/n) | 5/178 | 3/23 | 7/56 | 0.016 |
| Alcohol last day (y/n) | 37/146 | 1/25 | 10/53 | 0.11 |
| Coffee daily (y/n) | 131/52 | 21/8 | 46/20 | 0.95 |

### Clinical and neuropsychological assessment

Patients were diagnosed by trained clinicians using the Structured Clinical Interview for the DSM-IV, Axis I disorders (SCID-I) (21). Both patients and healthy controls were evaluated for IQ, using the Wechsler Abbreviated Scale of Intelligence (WASI) (22). Number of previous psychotic episodes, defined as an episode with score of ≥ 4 on Positive and Negative Syndrome Scale (PANSS) (23) items p1, p2, p3, p5, p6, or g9 for ≥ 1 week, was assessed. Current symptoms severity were evaluated using the PANSS, the Montgomery and Åsberg Depression Rating Scale (MADRS) (24), and the Young Mania Rating Scale (YMRS) (25). Current sleep disturbances were evaluated with the 4 sleep-related items of the Inventory of Depressive Symptoms (IDS) (26).

### Visual evoked potentials

The VEP modulation paradigm was adopted from Normann et al. (9), and all experimental procedures were performed exactly as described previously (27), with postintervention VEPs assessed at 120 s and 220 s (post 1), 380 s and 480 s (post 2), ∼30 min and ∼32 min (post 3), and ∼54 min and ∼56 min (post 4) after 10 minutes of checkerboard stimulation at a spatial frequency of 1 cycle/degree and a temporal frequency at 2 reversals per second (Fig. 1). To monitor constant fixation throughout the experiment, all participants focused on a fixation point at the centre of the screen and were asked to press a button when it changed color. EEG was recorded from a BioSemi ActiveTwo amplifier, with 64 Ag-AgCl sintered electrodes distributed across the scalp according to the international 10-20 system, and 4 electrodes located around the eyes to acquire horizontal and vertical electro-oculograms. Potentials at each channel were sampled at 2048 Hz with respect to a common mode sense, with a driven right leg electrode minimizing common mode voltages.

**Figure 1.**
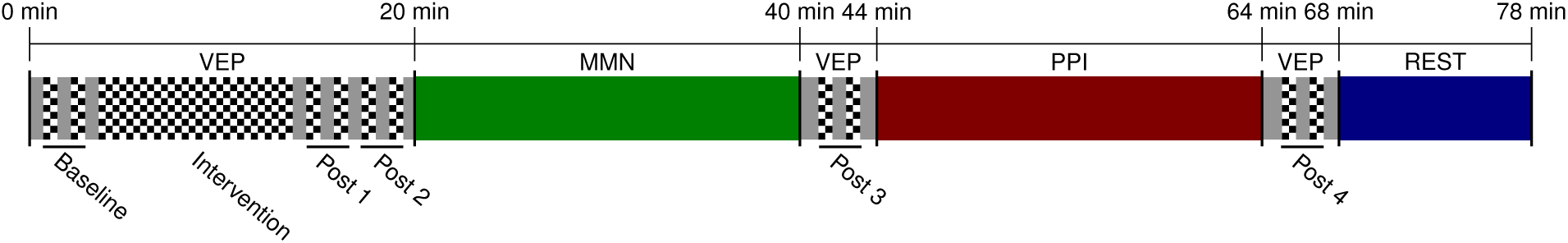
Experimental timeline. **VEP**: visual evoked potential paradigm, **MMN**: mismatch negativity paradigm, **PPI**: prepulse inhibition paradigm, **REST**: resting state EEG.

Signal processing was performed using MATLAB and the EEGLAB toolbox (28). Offline, recordings were downsampled to 512 Hz. Noisy channels were removed using PREP pipeline (29) with default settings, before re-referencing to remaining channel average, interpolation of removed channels, and finally a second re-referencing to the postinterpolation average. Data were band pass-filtered between 0.1 and 40 Hz. Event markers were adjusted to account for a latency of 20 ms in the visual presentation, measured with a BioSemi PIN diode, before epoch extraction at 200 ms pre- to 500 ms post-stimulus, and subsequent baseline correction. Artifactual muscle, eye blink, and eye movement components were removed with SASICA (30) using default parameters after independent component analysis using the SOBI algorithm. Epochs with a drift exceeding 100 μV were removed, and channels were rereferenced to the AFz electrode.

VEPs were averaged according to subject and pairs of blocks (baseline, post 1, post 2, post 3 and post 4), and components C1, P1, and N1b were extracted from the Oz electrode as the minimum amplitude between 50-100 ms post-stimulus, maximum amplitude between 80-140 ms, and mean amplitude between the first negative and halfway to the first positive peak after P1 (∼150-190 ms post-stimulus), respectively.

### Statistical analysis

Statistical analysis was performed in R version 3.6.0 (31). For all analyses except sensitivity analyses for separate diagnoses, patients with SZ, schizophreniform disorder, and schizoaffective disorder were considered conjointly, as were patients with BDI, BDII, BD NOS, and cyclothymia, with the resulting groups being referred to as SZ spectrum disorders (n=31) and BD spectrum disorders (n=69), respectively.

Participants with outlying difference scores (baseline amplitudes subtracted from postintervention amplitudes) for a VEP component (C1, P1, or N1b) at one or more postintervention assessments had all their postintervention assessments excluded from analysis for that particular component. Outliers were identified according to the median absolute deviation procedure implemented in R package `Routliers` (32), with 3 median absolute deviations as threshold, yielding 13 outliers for N1b modulation (SZ spectrum: 1, BD spectrum: 2, HC: 10). This procedure ensured a normal distribution of linear model residuals.

All analyses, except tests of baseline VEP component amplitudes, were performed directly on difference scores (baseline amplitudes subtracted from postintervention assessment amplitudes). Linear models were evaluated with type-II analyses of deviance implemented in R package `car` (33) to yield unbiased estimates of χ^2^ along with p-values for each predictor, or t-scores for intercepts. Outcomes that were not changing over time were assessed with two-tailed t-tests. All p-values are reported in uncorrected form, whereas significance thresholds were adjusted according to the effective number of independent comparisons within sets of analyses, by Sidak (34) or Bonferroni correction for continuous or categorical variables, respectively. This procedure yielded an α = 0.020 for primary analyses, i.e. modulation of C1, P1, and N1b modelled by diagnosis, time, and diagnosis x time, and for secondary analyses: i) α = 0.016 in the pairwise comparisons of N1b modulation between diagnoses, ii) α = 0.014 in the models of N1b modulation with clinical variables as predictors, and iii) α = 0.010 in the models of N1b modulation with groups of psychotropic medications as predictors. Lastly, the series of sensitivity tests, that were performed to examine the robustness of primary or secondary results, inherited significance thresholds from their parent analysis.

## Results

### N1b modulation is reduced in SZ and BD spectrum disorders

In this sample, there was modulation of VEP components C1 (t = 6.6, p = 7.3 x 10^−11^), P1 (t = 5.2, p = 2.9 x 10^−7^), and N1b (t = -7.7, p = 4.1 x 10^−14^) after prolonged visual stimulation (Figs. 2B-H). Further, there was no significant association between diagnosis and baseline amplitudes of either component C1 (χ^2^ = 3.4, p = 0.18), P1 (χ^2^ = 1.1, p = 0.54), or N1b (χ^2^ = 4.56, p = 0.10) (Fig. 2A).

**Figure 2.**
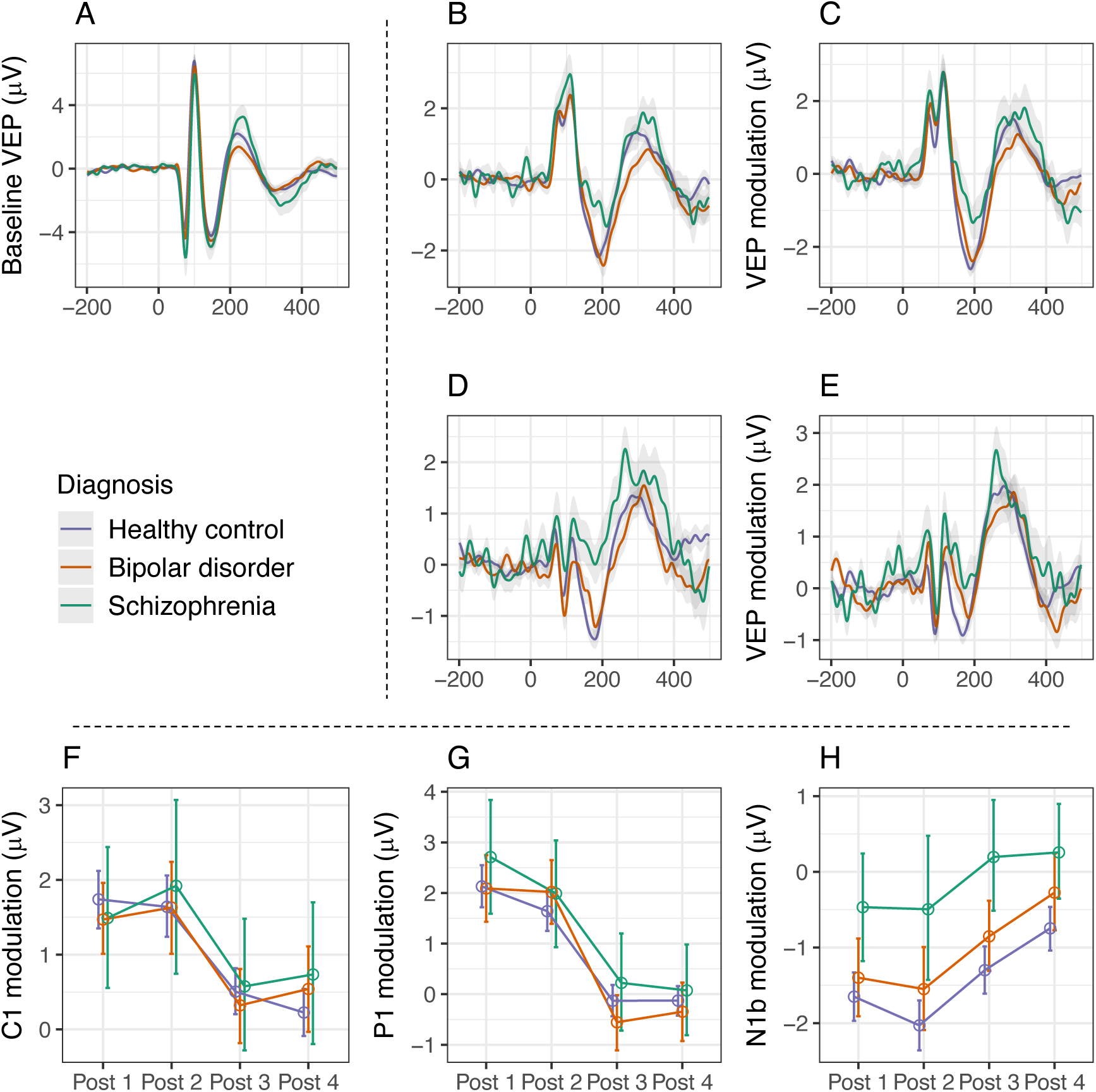
**A.** Visual evoked potentials (VEP) at baseline, by diagnostic group. VEPs were measured at the occiput (Oz), with anterior reference (AFz). **B.** VEP modulation (baseline VEP subtracted from postintervention VEP) at postintervention assessment 1 (2-4 min after prolonged visual stimulation), by diagnostic group. **C.** VEP modulation at postintervention assessment 2 (6-8 min after prolonged visual stimulation), by diagnostic group. **D.** VEP modulation at postintervention assessment 3 (30-32 min after prolonged visual stimulation), by diagnostic group. **E.** VEP modulation at postintervention assessment 4 (54-56 min after prolonged visual stimulation), by diagnostic group. **F.** C1 modulation (baseline C1 amplitudes subtracted from postintervention C1 amplitudes) at postintervention assessments 1-4, by diagnostic group. No difference in C1 modulation was detected between diagnostic groups (χ^2^ = 0.5, p = 0.78). **G.** P1 modulation at postintervention assessments 1-4, by diagnostic group. No difference in P1 modulation was detected betweeen diagnostic groups (χ^2^ = 2.7, p = 0.25). **H.** N1b modulation at postintervention assessments 1-4, by diagnostic group. N1b modulation was significantly different between healthy controls, BD patients, and SZ spectrum patients (χ^2^ = 37.9, p = 5.9 x 10^−9^).

The general linear model for N1b modulation with diagnostic group (SZ spectrum vs BD spectrum vs HC), time (postintervention assessments 1 vs 2 vs 3 vs 4), and diagnostic group x time interaction as predictors revealed an effect of diagnostic group (χ^2^ = 37.9, p = 5.9 x 10^−9^) and time (χ^2^ = 51.1, p = 4.6 x 10^−11^) on modulation of VEP component N1b (Fig. 2H, Table 2, Supplementary fig. S1), with no interaction effect (χ^2^ = 1.4, p = 0.97), demonstrating that N1b modulation was different between the diagnostic groups, and that N1b modulation waned over time across diagnostic groups. Corresponding general linear models did not demonstrate differences between diagnostic groups in the modulation of components C1 (χ^2^ = 0.5, p = 0.78, Fig. 2F) or P1 (χ^2^ = 2.7, p = 0.25, Fig. 2G), and further analyses for these components were not pursued.

**Table 2.**
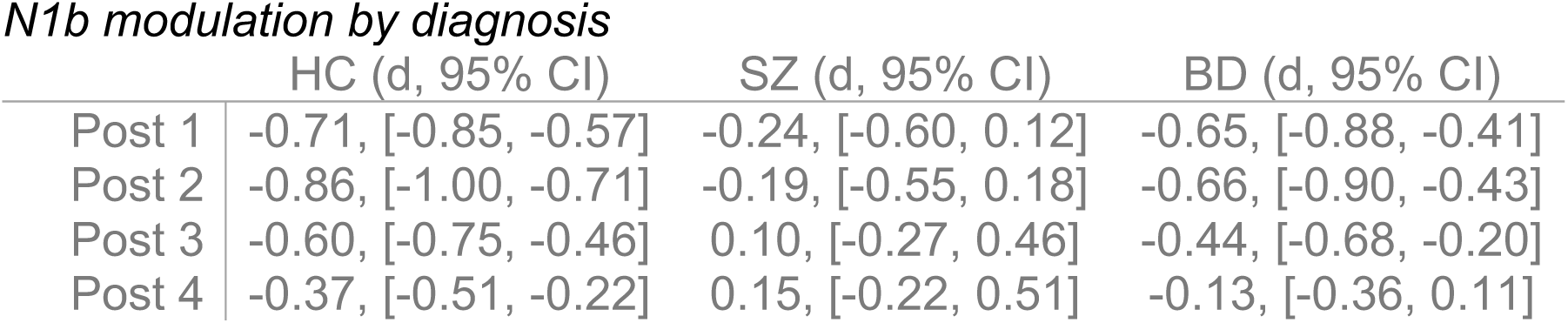
Modulation of VEP component N1b at post 1-4 assessments for participants with schizophrenia spectrum disorder (SZ), bipolar spectrum disorders (BD), or healthy controls (HC).

|  | HC (d, 95% CI) | SZ (d, 95% CI) | BD (d, 95% CI) |
| --- | --- | --- | --- |
| Post 1 | -0.71, [-0.85, -0.57] | -0.24, [-0.60, 0.12] | -0.65, [-0.88, -0.41] |
| Post 2 | -0.86, [-1.00, -0.71] | -0.19, [-0.55, 0.18] | -0.66, [-0.90, -0.43] |
| Post 3 | -0.60, [-0.75, -0.46] | 0.10, [-0.27, 0.46] | -0.44, [-0.68, -0.20] |
| Post 4 | -0.37, [-0.51, -0.22] | 0.15, [-0.22, 0.51] | -0.13, [-0.36, 0.11] |

Pairwise comparisons showed that modulation of N1b after prolonged visual stimulation was impaired in patients with SZ spectrum (χ^2^ = 35.1, p = 3.1 x 10^−9^) and in patients with BD spectrum disorders (χ^2^ = 7.0, p = 8.2 x 10^−3^) relative to controls. The impairment was more pronounced in SZ spectrum than in BD spectrum disorders (χ^2^ = 14.2, p = 1.7 x 10^−4^, Fig. 2H). Moreover, sensitivity analyses of separate diagnoses showed that N1b modulation was reduced in SZ alone (χ^2^ = 35.9, p = 2.1 x 10^−9^) and in BDI alone (χ^2^ = 6.4, p = 0.012), but not in BDII alone (χ^2^ = 2.2, p = 0.14).

We performed a series of sensitivity tests to examine the robustness of the effect of diagnosis on N1b modulation. The effect of diagnosis on N1b modulation remained significant when controlling for baseline amplitudes, sex, and age (χ^2^ = 28.5, p = 6.6 x 10^−7^), as well as when controlling for mood states, as indexed by MADRS and YMRS (χ^2^ = 19.6, p = 5.6 x 10^−5^), and when controlling for IQ (χ^2^ = 21.2, p = 2.5 x 10^−5^), current sleep disturbance (χ^2^ = 30.2, p = 2.8 x 10^−7^), and daily use of tobacco, monthly use of cannabis, or use of alcohol within the last day before examination (χ^2^ = 32.0, p = 1.1 x 10^−7^). Further, the effect of diagnosis on N1b modulation remained significant when considering only unmedicated patients (n = 20) against healthy controls (χ^2^ = 18.7, p = 8.7 x 10^−5^). The effect of diagnosis on N1b modulation also was significant in the model where ouliers were included (χ^2^ = 29.1, p = 4.8 x 10^−7^). Lastly, although due to an error in the gaming controller used for responses to on-screen dot color changes, these response data were missing for 40.4% of the current sample, there were no significant group differences in the proportion of correct responses to the on-screen dot color changes (t = -0.4, p = 0.71), indicating that the attention afforded to the prolonged visual stimulation did not differ between patients and controls.

### Associations between N1b modulation and clinical states

Within patients, decreased N1b modulation was significantly associated with greater symptom severity, as indexed with PANSS total (χ^2^ = 10.8, p = 1.0 x 10^−3^, Fig. 3A), and nominally with number of psychotic episodes (χ^2^ = 4.9, p = 0.027, Fig. 3B). We performed a series of sensitivity tests to examine whether these associations between psychotic illness severity and reduced N1b modulation remained within diagnostic spectra. The association between N1b modulation and PANSS total was not reproduced within the BD spectrum (χ^2^ = 0.3, p = 0.56), but did not remain significant within the SZ spectrum (χ^2^ = 3.3, p = 0.07). The association between N1b modulation and number of psychotic episodes remained only nominally significant within the BD spectrum (χ^2^ = 5.6, p = 0.018), and was absent within the SZ spectrum (χ^2^ = 0.1, p = 0.72). Finally, modulation of component N1b was unaffected by mood state as indexed either with MADRS (χ^2^ = 0.3, p = 0.61, Fig. 3C) or YMRS (χ^2^ = 0.0, p = 0.99, Fig. 3D).

**Figure 3.**
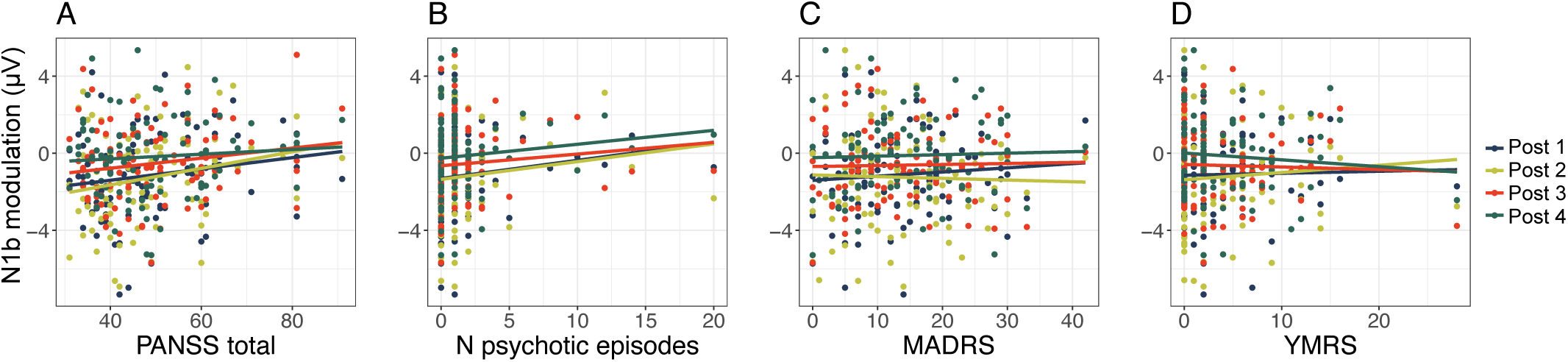
Associations between clinical states and N1b modulation in patients, at postintervention assessments 1-4. **A.** N1b modulation was associated with PANSS (Positive and Negative Syndrome Scale) total scores in patients (χ^2^ = 10.8, p = 1.0 x 10^−3^). **B.** N1b modualation showed a nominally significant association with number of psychotic episodes in patients (χ^2^ = 4.9, p = 0.027). **C.** N1b modulation was not associated with MADRS (Montgomery and Åsberg Depression Rating Scale) scores in patients (χ^2^ = 0.3, p = 0.61). **D.** N1b modulation was not associated with YMRS (Young Mania Rating Scale) scores in patients (χ^2^ = 0.0, p = 0.99).

### N1b modulation is further reduced in patients using antiepileptic or antipsychotic medication

Across all patients, N1b modulation was lower in users of antipsychotic medication (χ^2^ = 8.3, p = 3.9 x 10^−3^), with a similar trend observed in users of antiepileptic medication (χ^2^ = 3.5, p = 0.062), than in non-users. Thus, we performed a series of sensitivity tests to further examine the relationship between psychotropic medication use and N1b modulation. In BD patients, N1b modulation was more severely impaired among users of antiepileptics (χ^2^ = 9.3, p = 2.3 x 10^−3^, Fig. 4A), and nominally in users of antipsychotics (χ^2^ = 4.4, p = 0.035, Fig. 4B). Further, the association in BD patients between lamotrigine and N1b modulation (χ^2^ = 6.8, p = 8.9 x 10^−3^) was comparable to the effect of antiepileptics in general. Within BDII patients only, N1b modulation was still lower among users of antiepileptics (χ^2^ = 9.1, p = 2.5 x 10^−3^, n = 11), and nominally lower among users of antipsychotics (χ^2^ = 4.7, p = 0.03, n = 7). However, within BDI patients only, N1b modulation did not remain significantly lowered among users of antiepileptics (χ^2^ = 3.2, p = 0.07, n = 11), nor among users of antipsychotics (χ^2^ = 0.6, p = 0.45, n = 17). There was no evidence for lowered N1b modulation among antipsychotics users with SZ (χ^2^ = 0.1, p = 0.70, Fig. 4C), and only one SZ patient used antiepileptics. Further, when controlling for psychotic symptom severity and diagnosis, the association with reduced N1b modulation remained for antiepileptics use (χ^2^ = 11.4, p = 7.1 x 10^−4^), but not for antipsychotics use (χ^2^ = 2.4, p = 0.12). Finally, there was no evidence for any change in N1b modulation among patients using either lithium (χ^2^ = 0.6, p = 0.43), antidepressants (χ^2^ = 1.3, p = 0.25), or anxiolytics/hypnotics (χ^2^ = 1.5, p = 0.22).

**Figure 4.**
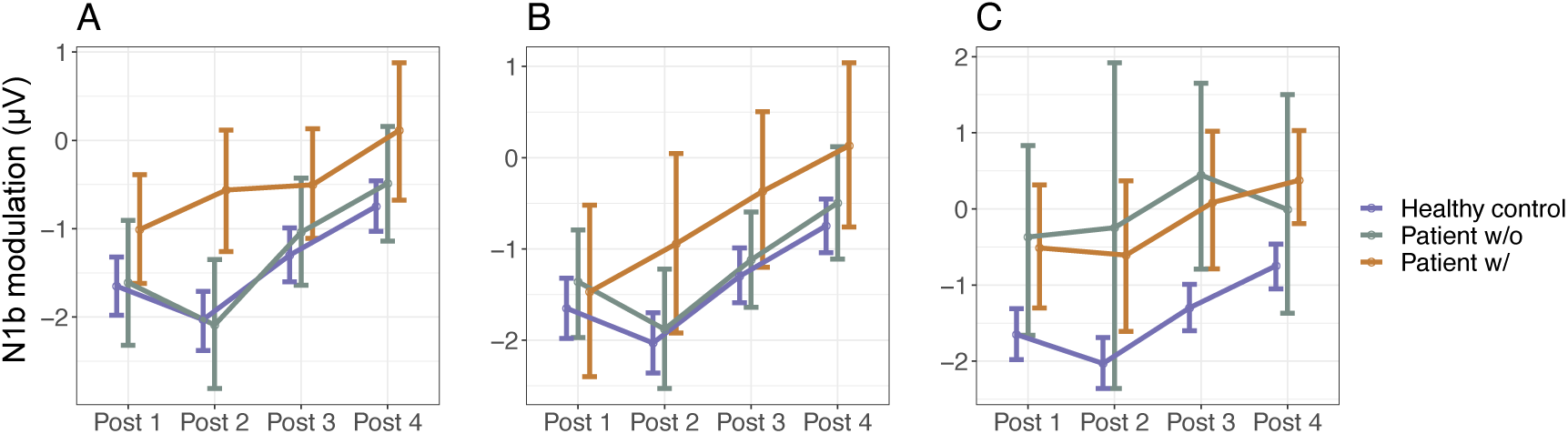
**A.** Antiepileptics use in patients with BD. N1b modulation at postintervention assessments 1-4 was lower in BD patients using antiepileptics (w/) than in BD patients not using antiepileptics (w/o) (χ^2^ = 9.3, p = 2.3 x 10^−3^). N1b modulation for healthy controls is represented for comparison. **B.** Αntipsychotics use in patients with BD. N1b modulation at postintervention assessments 1-4 was tendentially lower in BD patients using antipsychotics (w/) than in BD patients not using antipsychotics (w/o) (χ^2^ = 4.4, p = 0.035). **C.** Antipsychotics use in patients with SZ spectrum disorders. N1b modulation at postintervention assessments 1-4 was not significantly different between SZ spectrum patients using antipsychotics (w/) and SZ spectrum patients not using antipsychotics (w/o) (χ^2^ = 0.1, p = 0.70).

## Discussion

This study of LTP-like visual cortical plasticity in patients with SZ, BDI, and BDII, and healthy controls, yielded four main results. First, relative to age-matched healthy controls, modulation of the N1b component of the VEP after prolonged visual stimulation was significantly reduced in SZ and BDI, but not BDII patients, with stronger reductions seen in patients with more severe psychotic symptoms. Second, we observed significant association between N1b modulation and current use of medication, suggesting that N1b modulation is lower in patients using antiepileptic or antipsychotic medication. Third, we did not observe any significant associations between N1b modulation and mood states. Finally, we did not observe any significant difference between patients with BD or SZ spectrum disorders and healthy controls in the modulation of VEP components C1 or P1.

Modulation of VEP components in general, and the N1b component in particular, has been implicated as a candidate index of NMDAR-dependent LTP-like plasticity in the visual cortex. In humans, N1b modulation is dependent on high frequency or prolonged visual stimulation (10), and seems to have more robust response and a time-course more compatible with LTP than the modulation of other VEP components (27). Further, N1b component modulation after visual stimulation is orientation- and spatial frequency specific in humans, indicating a synapse specificity of N1b modulation similar to LTP (35,36). The result that N1b component modulation after visual stimulation is reduced in SZ and BDI patients is therefore in line with previous genetic (37,38), molecular (39), pharmacological (6), and anatomical evidence (40,41), strengthening the previously suggested hypothesis that NMDAR-dependent synaptic plasticity is affected in these psychiatric disorders (8).

The present results provide a demonstration of reduced N1b modulation in patients with BDI, an association which has not been examined previously. Further, the present results provide a clear demonstration of reduced N1b modulation in patients with SZ. Previously, two separate studies have compared VEP modulation between SZ patients and healthy controls, with the first study showing evidence for reduced N1b modulation in SZ (17), whereas the second study found no evidence for altered VEP modulation in SZ (18). In the former study (17), modulation of component C1 was also decreased in schizophrenia patients, albeit with lower certainty than for the N1b component. Rather than a checkerboard stimulus, the latter study used grating stimulus, which is well suited for manipulating stimulus orientation and assessing the input specificity of modulation effects, along with a higher frequency and shorter duration of the intervention as compared to the present and other studies (9,16). One or more of these conditions may have contributed to lower effect sizes and accordingly lower power in detecting group differences (18). Previously, two studies have compared VEP modulation between BDII patients and healthy controls, both observing tendencies of reduced modulation of N1, a component that is highly correlated with N1b (27) in BDII patients, comparable to the tendency of reduced N1b modulation observed in the present sample. Taken together, these converging results suggest that N1b modulation after prolonged visual stimulation, likely indexing NMDAR-dependent synaptic plasticity in the visual cortex (12-14), is reduced in SZ and BDI, but to a lesser extent in BDII.

Further, we observed that reduction in N1b modulation was associated with higher psychotic symptom severity, as indexed with PANSS total and, nominally, as indexed with number of psychotic episodes, but not with mood states, as indexed with MADRS or YMRS. The association between N1b modulation and psychotic symptom severity was, however, driven to a large degree by diagnosis, and did not remain significant within diagnostic groups after controlling for multiple comparisons, although tendencies were preserved in the associations with PANSS total within SZ patients and with number of psychotic episodes within BD patients. Nevertheless, since N1b modulation is more reduced among patients with higher PANSS total scores and, nominally, among patients with a history of more psychotic episodes, and since N1b modulation is more reduced in disorders defined by psychosis or with higher prevalence of psychosis, there is reason to suggest that psychotic symptoms in particular are related to reduced N1b modulation.

Reduced N1b modulation was also associated with use of psychotropic medication among patients. First, use of antiepileptics, particularly lamotrigine, was associated with reduced N1b modulation in BD patients. Further, the association between antiepileptics use and reduced N1b modulation remained when controlling for specific diagnosis and for PANSS total scores. Thus, the decreased N1b modulation among antiepileptics users is likely not explained by diagnosis or by psychotic symptom severity. Although the extent to which antiepileptics decrease the probability of NMDAR-dependent synaptic plasticity remains to be clarified, antiepileptics promote GABAA-mediated inhibition, increase sodium channel resistance, inhibit glutamate release (42,43), and have been shown to decrease LTP in hippocampal slices (44). Lamotrigine likely inhibits glutamate release through increasing sodium channel resistance, which could potentially contribute to the reduced N1b modulation. However, future pharmacological studies using a randomized controlled design would be needed to carefully test this hypothesis. Second, antipsychotics use was associated with reduced N1b modulation among BD patients, but not among schizophrenia patients. However, the association between N1b modulation and antipsychotics use did not remain after controlling for diagnosis and psychotic symptom severity, suggesting that this association could reflect intrinsic differences in antipsychotics users vs non-users, rather than a direct effect of antipsychotics use on LTP-like plasticity.

## Conclusion

The present study demonstrated impaired LTP-like plasticity in patients with SZ and BDI, but not in patients with BDII. Together with previous genetic, pharmacological, and anatomical research, these results implicate aberrant synaptic plasticity as a pathophysiological mechanism in SZ and BD.

## Acknowledgments

This study was funded by the Research Council of Norway, the South-Eastern Norway Regional Health Authority, Oslo University Hospital and a research grant from Mrs. Throne-Holst. The authors report no biomedical financial interests or potential conflicts of interest.

## Supplementary Figure

**Supplementary Figure 1.**
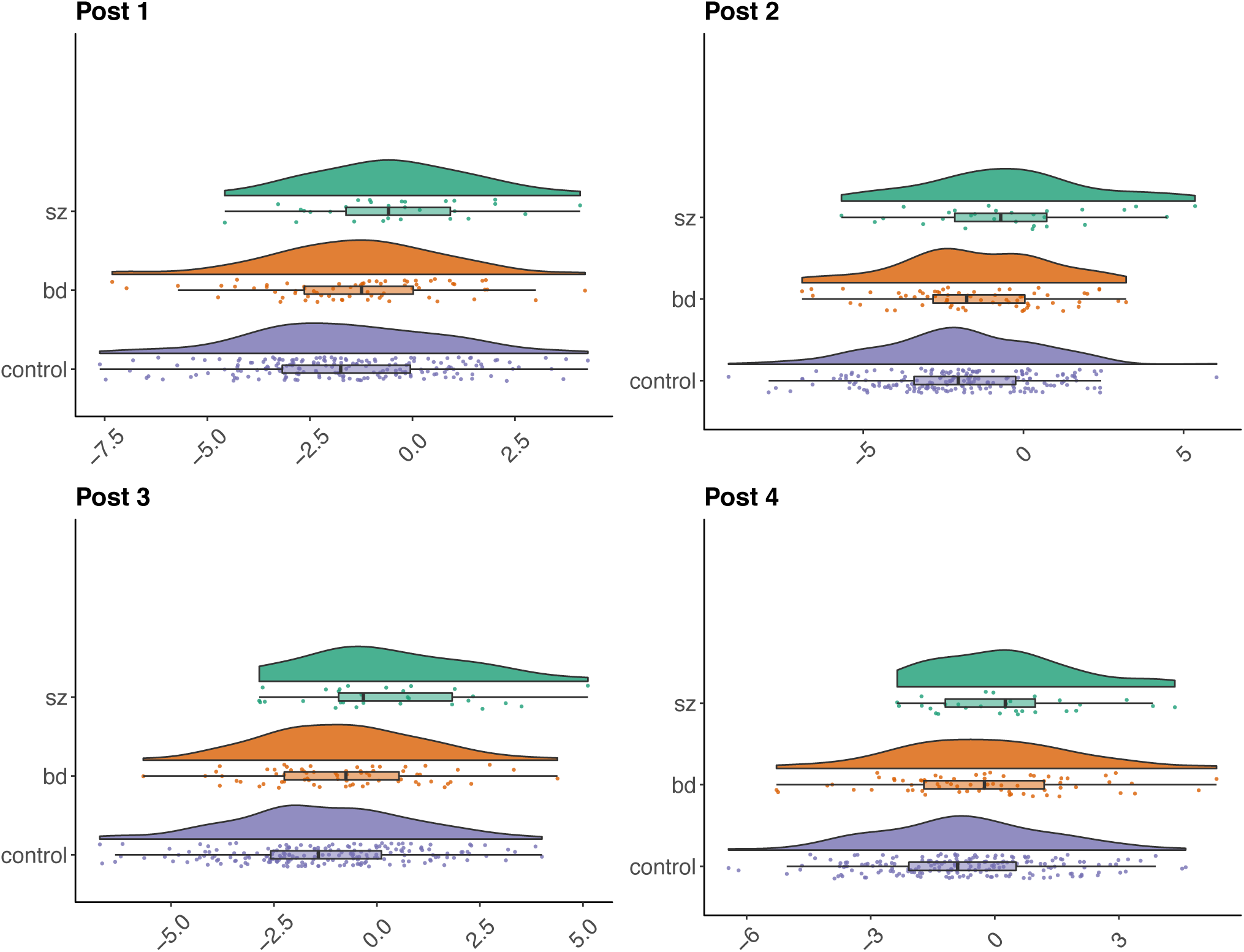
Raincloud plots of N1b modulation at post 1-4 assessments for healthy controls, patients with bipolar spectrum disorder, or patients with schizophrenia spectrum disorders.

## References

1. McGrath J, Saha S, Chant D, Welham J (2008): Schizophrenia: A concise overview of incidence, prevalence, and mortality. Epidemiol Rev. 30: 67–76.

2. Merikangas KR, Jin R, He J-P, Kessler RC, Lee S, Sampson NA, et al. (2011): Prevalence and correlates of bipolar spectrum disorder in the world mental health survey initiative. Arch Gen Psychiatry. 68: 241–251.

3. Dell’Aglio JC Jr, Basso LA, Argimon II de L, Arteche A (2013): Systematic review of the prevalence of bipolar disorder and bipolar spectrum disorders in population-based studies. Trends in Psychiatry and Psychotherapy. 35: 99–105.

4. Omdal R, Brokstad K, Waterloo K, Koldingsnes W, Jonsson R, Mellgren SI (2005): Neuropsychiatric disturbances in SLE are associated with antibodies against NMDA receptors. European Journal of Neurology. 1–7.

5. Dalmau J, Lancaster E, Martinez-Hernandez E, Rosenfeld MR, Balice-Gordon R (2011): Clinical experience and laboratory investigations in patients with anti-NMDAR encephalitis. Lancet Neurol. 10: 63–74.

6. Javitt DC, Zukin SR (1991): Recent advances in the phencyclidine model of schizophrenia. American Journal of Psychiatry. 148: 1301–1308.

7. Krystal JH, Karper LP, Seibyl JP, Freeman GK, Delaney R, Bremner JD, et al. (1994): Subanesthetic effects of the noncompetitive NMDA antagonist, ketamine, in humans: Psychotomimetic, perceptual, cognitive, and neuroendocrine responses. JAMA Psychiatry. 51: 199–214.

8. Stephan KE, Baldeweg T, Friston KJ (2006): Synaptic plasticity and dysconnection in schizophrenia. Biological Psychiatry. 59: 929–939.

9. Normann C, Schmitz D, Fürmaier A, Döing C, Bach M (2007): Long-term plasticity of visually evoked potentials in humans is altered in major depression. Biol Psychiatry. 62: 373–380.

10. Teyler TJ, Hamm JP, Clapp WC, Johnson BW, Corballis MC, Kirk IJ (2005): Long-term potentiation of human visual evoked responses. European Journal of Neuroscience. 21: 2045–2050.

11. Clapp WC, Hamm JP, Kirk IJ, Teyler TJ (2012): Translating long-term potentiation from animals to humans: A novel method for noninvasive assessment of cortical plasticity. Biol Psychiatry. 71: 496–502.

12. Frenkel MY, Sawtell NB, Diogo ACM, Yoon B, Neve RL, Bear MF (2006): Instructive effect of visual experience in mouse visual cortex. Neuron. 51: 339–349.

13. Cooke SF, Bear MF (2010): Visual experience induces long-term potentiation in the primary visual cortex. J Neurosci. 30: 16304–16313.

14. Cooke SF, Bear MF (2012): Stimulus-selective response plasticity in the visual cortex: An assay for the assessment of pathophysiology and treatment of cognitive impairment associated with psychiatric Disorders. Biol Psychiatry. 71: 487–495.

15. Zak N, Moberget T, Bøen E, Boye B, Waage TR, Dietrichs E, et al. (2018): Longitudinal and cross-sectional investigations of long-term potentiation-like cortical plasticity in bipolar disorder type II and healthy individuals. Transl Psychiatry. 8: 103.

16. Elvsåshagen T, Moberget T, Bøen E, Boye B, Englin NOA, Pedersen PØ, et al. (2012): Evidence for impaired neocortical synaptic plasticity in bipolar II disorder. Biol Psychiatry. 71: 68–74.

17. Çavuş I, Reinhart RMG, Roach BJ, Gueorguieva R, Teyler TJ, Clapp WC, et al. (2012): Impaired visual cortical plasticity in schizophrenia. Biol Psychiatry. 71: 512–520.

18. Wynn JK, Roach BJ, McCleery A, Marder SR, Mathalon DH, Green MF (2019): Evaluating visual neuroplasticity with EEG in schizophrenia outpatients. Schizophrenia Research. doi: 10.1016/j.schres.2019.08.015.

19. Ho DE, Imai K, King G, Stuart EA (2011): Nonparametric preprocessing for parametric causal inference. Journal of Statistical Software. 42: 1–28.

20. Richard G, Kolskår K, Sanders A-M, Kaufmann T, Petersen A, Doan NT, et al. (2018): Assessing distinct patterns of cognitive aging using tissue-specific brain age prediction based on diffusion tensor imaging and brain morphometry. PeerJ. 6: e5908.

21. First MB, Spitzer RL, Gibbon M, Williams J (1996): Structured Clinical Interview for DSM-IV Axis I Disorders, Clinician Version (SCID-CV). American Psychiatric Press.

22. Wechsler D (2007): Wechsler Abbreviated Scale of Intelligence (WASI). Stockholm: Pearson Assessment.

23. Kay SR, Fiszbein A, Opler LA (1987): The Positive and Negative Syndrome Scale (PANSS) for Schizophrenia. Schizophrenia Bulletin. 13: 261–276.

24. Montgomery SA, Åsberg M (1979): A new depression scale designed to be sensitive to change. Br J Psychiatry. 134: 382–389.

25. Young RC, Biggs JT, Ziegler VE, Meyer DA (1978): A rating scale for mania: Reliability, validity and sensitivity. Br J Psychiatry. 133: 429–435.

26. Trivedi MH, Rush AJ, Ibrahim HM, Carmody TJ, Biggs MM, Suppes T, et al. (2004): The Inventory of Depressive Symptomatology, Clinician Rating (IDS-C) and Self-Report (IDS-SR), and the Quick Inventory of Depressive Symptomatology, Clinician Rating (QIDS-C) and Self-Report (QIDS-SR) in public sector patients with mood disorders: a psychometric evaluation. Psychol Med. 34: 73–82.

27. Valstad M, Moberget T, Roelfs D, Slapø NB, Timpe CMF, Beck D, et al. (2020): Experience-dependent modulation of the visual evoked potential: testing effect sizes, retention over time, and associations with age in 415 healthy individuals. bioRxiv. 2020.01.27.916692.

28. Delorme A, Makeig S (2004): EEGLAB: an open source toolbox for analysis of single-trial EEG dynamics including independent component analysis. Journal of Neuroscience Methods. 134: 9–21.

29. Bigdely-Shamlo N, Mullen T, Kothe C, Su K-M, Robbins KA (2015): The PREP pipeline: standardized preprocessing for large-scale EEG analysis. Front Neuroinform. 9: B153.

30. Chaumon M, Bishop DVM, Busch NA (2015): A practical guide to the selection of independent components of the electroencephalogram for artifact correction. Journal of Neuroscience Methods. 250: 47–63.

31. R Core Team (2019): R: A Language and Environment for Statistical Computing. 3rd ed.

32. Delacre M, Klein O (2019): Routliers: robust outliers detection. R package version 0.0.0.3.

33. Fox J, Weisberg S (2019): An {R} companion to applied regression.

34. Li J, Ji L (2005): Adjusting multiple testing in multilocus analyses using the eigenvalues of a correlation matrix. Heredity 2005 95:3. 95: 221–227.

35. McNair NA, Clapp WC, Hamm JP, Teyler TJ, Corballis MC, Kirk IJ (2006): Spatial frequency-specific potentiation of human visual-evoked potentials. NeuroReport. 17: 739–741.

36. Ross RM, McNair NA, Fairhall SL, Clapp WC, Hamm JP, Teyler TJ, Kirk IJ (2008): Induction of orientation-specific LTP-like changes in human visual evoked potentials by rapid sensory stimulation. Brain Research Bulletin. 76: 97–101.

37. Schizophrenia Working Group of the Psychiatric Genomics Consortium (2014): Biological insights from 108 schizophrenia-associated genetic loci. Nature. 511: 421–427.

38. Stahl EA, Breen G, Forstner AJ, McQuillin A, Ripke S, Trubetskoy V, et al. (2019): Genome-wide association study identifies 30 loci associated with bipolar disorder. Nat Genet. 51: 793–803.

39. Akbarian S, Sucher NJ, Bradley D, Tafazzoli A, Trinh D, Hetrick WP, et al. (2003): Selective alterations in gene expression for NMDA receptor subunits in prefrontal cortex of schizophrenics. 1–12.

40. Garey L, Ong W, Patel T, Kanani M, Davis A, Mortimer A, et al. (1998): Reduced dendritic spine density on cerebral cortical pyramidal neurons in schizophrenia. Journal of Neurology, Neurosurgery & Psychiatry. 65: 446–453.

41. Black JE, Kodish IM, Grossman AW, Klintsova AY, Orlovskaya D, Vostrikov V, et al. (2004): Pathology of Layer V pyramidal neurons in the prefrontal cortex of patients with schizophrenia. American Journal of Psychiatry. doi: 10.1176/appi.ajp.161.4.742.

42. Rogawski MA, Löscher W (2004): The neurobiology of antiepileptic drugs. Nature Reviews Neuroscience. 5: 553–564.

43. Ikonomidou C, Turski L (2010): Antiepileptic drugs and brain development. Epilepsy Research. 88: 11–22.

44. Lee GY-P, Brown LM, Teyler TJ (1996): The effects of anticonvulsant drugs on long-term potentiation (LTP) in the rat hippocampus. Brain Research Bulletin. 39: 39–42.

